# Enhanced cellular longevity arising from environmental fluctuations

**DOI:** 10.1101/2023.07.05.547867

**Authors:** Yuting Liu, Zhen Zhou, Songlin Wu, Gavin Ni, Alex Zhang, Lev S. Tsimring, Jeff Hasty, Nan Hao

## Abstract

Cellular longevity is regulated by both genetic and environmental factors. However, the interactions of these factors in the context of aging remain largely unclear. Here, we formulate a mathematical model for dynamic glucose modulation of a core gene circuit in yeast aging, which not only guided the design of pro-longevity interventions, but also revealed the theoretical principles underlying these interventions. We introduce the dynamical systems theory to capture two general means for promoting longevity - the creation of a stable fixed point in the “healthy” state of the cell and the dynamic stabilization of the system around this healthy state through environmental oscillations. Guided by the model, we investigate how both of these can be experimentally realized by dynamically modulating environmental glucose levels. The results establish a paradigm for theoretically analyzing the trajectories and perturbations of aging that can be generalized to aging processes in diverse cell types and organisms.

## Introduction

Aging is driven by accumulation of cellular and genetic damage resulting from intertwined biological processes that are intrinsic to the individual and which are influenced by environmental factors (Lopez-Otin et al., 2013; Mahmoudi and Brunet, 2012; McMurray and Gottschling, 2004; Melzer et al., 2020; Vijg and Suh, 2013). Novel approaches to reduce global healthcare burdens of chronic diseases and aging ultimately demand increased understanding of aging biology and the interactions of the pillars of aging that include diverse yet deeply linked factors such as epigenetics, stress, metabolism, and others (Kennedy et al., 2014; Lopez-Otin et al., 2013). Previous studies in model organisms have been focused on measuring lifespan as a static endpoint assay and have identified many genes, the deletion or overexpression of which affects the lifespan (Fontana et al., 2010; Guarente and Kenyon, 2000; Kaeberlein and Kennedy, 2005; Kenyon, 2010; Kuningas et al., 2008; McCormick et al., 2015). An emerging challenge is to understand how these genes interact with one another and operate collectively to drive the aging processes and determine the final lifespan. Because of the intricacies of aging-related processes, traditional reductionist approaches cannot address the totality of such complexity. Instead, new systems-level approaches that integrate stochastic and nonlinear dynamic models with large time-trace data sets are required.

To this end, we set out to quantify and model replicative aging of the budding yeast *Saccharomyces cerevisiae*, a genetically tractable model for aging of mitotic cell types in mammals, such as stem cells. Using microfluidics coupled with time-lapse microscopy (O’Laughlin et al., 2020), we quantitatively tracked the aging processes in a large number of single yeast cells and found that isogenic cells age with two different types of phenotypic changes (Jin et al., 2019; Li et al., 2020; Li et al., 2017; Paxman et al., 2022): about half of cells continuously produced daughters with an elongated morphology during later stages of life (designated as “Mode 1” aging). In contrast, the other half produced small, round daughter cells until death (designated as “Mode 2” aging). Mode 1 aging is driven by ribosomal DNA (rDNA) silencing loss, resulting in dramatically enlarged and fragmented nucleoli, indicating nucleolar decline (Sinclair et al., 1997). In contrast, Mode 2 aging is driven by heme depletion and mitochondrial deterioration (Hughes and Gottschling, 2012).

In yeast, the lysine deacetylase Sir2 mediates chromatin silencing at rDNA to maintain the stability of this fragile genomic locus and the integrity of the nucleolus (Gartenberg and Smith, 2016; Kaeberlein et al., 1999; Saka et al., 2013; Sinclair and Guarente, 1997). The heme-activated protein (HAP) complex regulates the expression of genes important for heme biogenesis and mitochondrial function (Buschlen et al., 2003). We previously identified a mutual inhibition circuit of Sir2 and HAP that resembles a toggle switch to mediate the fate decision and divergent progression toward Mode 1 vs Mode 2 aging in single cells (Li et al., 2020). Guided by mathematical modeling of the endogenous system, we genetically engineered the Sir2-HAP circuit to reprogram aging trajectories and promote longevity (Li et al., 2020; Zhou et al., 2023). These studies revealed the design principles of genetic circuits for promoting cellular longevity under a static environmental condition. However, how environmental fluctuations impact the dynamical behaviors of genetic circuits to regulate the aging process remains largely unclear.

In this study, we used mathematical modeling approach to explore the possibility of rationally reprogramming aging by dynamic environmental inputs. Aging is strongly related to energy metabolism (Bartke et al., 2021; Roberts and Rosenberg, 2006). A growing number of studies revealed that metabolic alterations are a major hallmark of aging, whereas modulating energy metabolism by environmental factors can dramatically influence the phenotypes and rates of aging (Azzu and Valencak, 2017; Jazwinski, 2002; Petr et al., 2021; Ravera et al., 2019). For example, caloric restriction (CR) can promote longevity from yeast to mammals (Al-Regaiey, 2016; Arslan-Ergul et al., 2013; Fontana and Partridge, 2015; Hwangbo et al., 2020; Li et al., 2011; Liang et al., 2018; Taormina and Mirisola, 2014). Yet, a systematic, quantitative analysis of the impact of environmental factors on the complex networks of aging remain largely missing, making rational reprogramming of aging a challenging task. To this end, we developed a mathematical model to understand how environmental glucose alterations influence the dynamical behaviors of the core Sir2-HAP circuit and cellular longevity. Based on the single-cell data and dynamical systems theory, we identified two general approaches to extend lifespan by dynamically adjusting environmental glucose inputs - establishing a subtle balance (a stable fixed point) to stabilize the healthy state of the cell and driving dynamic stabilization of the system around this healthy state. Our model not only provides valuable biological insights for designing strategies to promote longevity but also uncovers the underlying theoretical principles behind these strategies, with broad applications across different cell types and organisms.

## Results

### Metabolic divergence in isogenic aging cells

To determine how aging affects cellular energy production under standard yeast growth conditions, we monitored cellular ATP levels throughout the lifespans of single cells using a genetically-encoded fluorescent biosensor (Lobas et al., 2019). We observed a gradual decline in cellular ATP levels in Mode 2, but not Mode 1 aging cells (Fig. 1A). To determine the metabolic processes underlying this difference in ATP production among aging cells, we monitored the levels of metabolic factors involved in energy production, such as glycolysis and respiration, and compared their dynamics between Mode 1 and Mode 2 cells.

**Fig 1.**
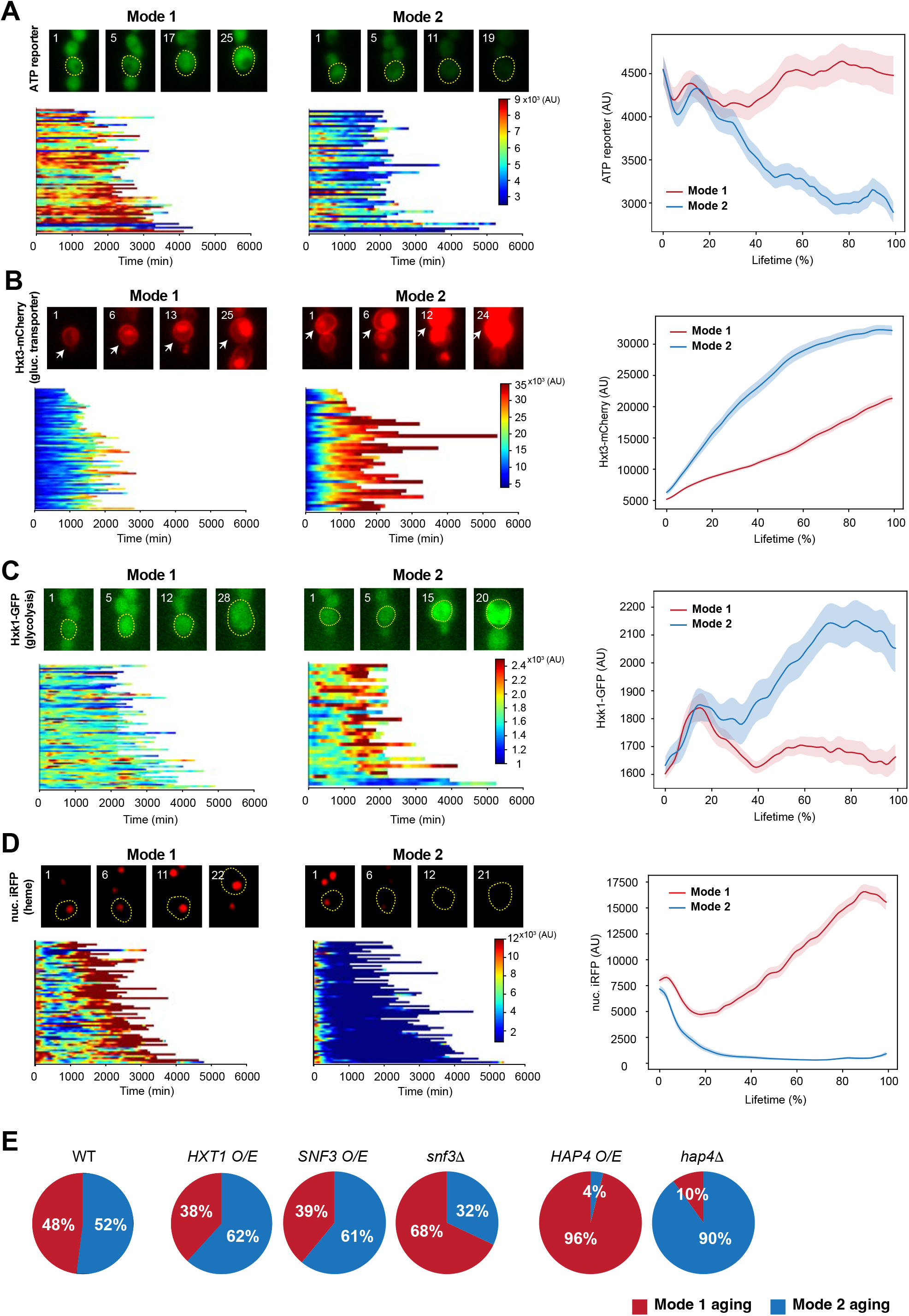
Distinct metabolic changes in Mode 1 vs Mode 2 aging. (**A-D**) Dynamics of an ATP reporter (A, n=116), Hxt3-mCherry (B, n=119), Hxk1-GFP (C, n=115), nuc. iRFP (D, n = 120) in Mode 1 and Mode 2 cells. (Top left of each panel) Representative time-lapse images of single Mode 1 and Mode 2 cells. Mother cells are circled in yellow, and the replicative age of the mother cell is shown at the top left corner of each image. (Bottom left of each panel) Single-cell color map trajectories in Mode 1 and Mode 2 cells. Each row represents the time trace of a single cell throughout its lifespan. Color represents the fluorescence intensity as indicated in the color bar. (Right of each panel) Average fluorescence time traces throughout the life spans in Mode 1 and Mode 2 cells. (**E**) Effects of metabolic gene perturbations on the fate decision in yeast aging. Pie charts show the percentages of Mode 1 (red) and Mode 2 (blue) in WT and mutants tested. O/E: overexpression.

We first examined changes in glycolysis during aging by tracking the expression of glucose transporters (Ozcan and Johnston, 1999), a glycolytic enzyme (Belinchon and Gancedo, 2007) and the fluorescence of a fructose-1,6-phosphate (FBP) glycolytic flux biosensor (Monteiro et al., 2019). The three glucose transporters we evaluated (Hxt1, Hxt2, and Hxt3) all showed dramatically increased expression in Mode 2 aging, but only modest changes in Mode 1 aging (Fig. 1B; Fig. S1, A and B), indicating that a marked elevation in glycolysis occurs specifically during Mode 2 aging. Consistent with this observation, the expression of the glycolytic enzyme Hxk1 also increased in Mode 2 but not in Mode 1 aging (Fig. 1C). Moreover, Mode 2 aging cells showed a more dramatic increase in the fluorescence of the glycolytic flux biosensor than Mode 1 cells (Fig. S1C). Heme abundance is an indicator of mitochondrial biogenesis and cellular respiration (Buschlen et al., 2003; Zhang et al., 2017). Using a nuclear-anchored infrared fluorescent protein (nuc. iRFP) (Filonov et al., 2011; Li et al., 2020), we tracked heme abundance in single aging cells and observed that it gradually elevated in Mode 1 aging, but dropped sharply in Mode 2 aging (Fig. 1D).

Together, these results indicated that Mode 1 aging features an age-induced transition from fermentation to respiration, in agreement with a previous report (Leupold et al., 2019), whereas Mode 2 aging elicits enhanced glycolysis but suppression of respiration, resulting in a decline in ATP production.

We next asked if the observed metabolic changes contribute to driving the divergence in Mode 1 and Mode 2 aging or whether they are simply the consequences of different aging routes. We genetically perturbed the glycolysis or respiration pathway and examined the effects on the fate decision in an isogenic aging population (Fig. 1E). We observed that overexpression of the glucose transporter Hxt1 or the glucose sensor Snf3, both of which elevate glucose uptake and glycolysis (Ozcan and Johnston, 1999), promoted Mode 2 aging, whereas deletion of Snf3 promoted Mode 1 aging. In addition, overexpression of Hap4, which is a major component of the HAP complex – the master transcriptional activator of yeast respiration (Forsburg and Guarente, 1989; Zhang et al., 2017), largely promoted Mode 1 aging, whereas deletion of Hap4 promoted Mode 2 aging, as previously reported (Li et al., 2020). These results demonstrate that metabolic alterations actively regulate the fate decision of single-cell aging – enhancing respiration or reducing glycolysis promotes Mode 1 aging, whereas elevating glycolysis or reducing respiration promotes Mode 2 aging.

### An optimal level of caloric restriction (CR) for lifespan extension

Environmental glucose levels can modulate metabolic processes, such as glycolysis and respiration (Johnston, 1999; Rolland et al., 2002). To systematically determine how alterations in glucose levels influence the fate decision in aging and the cellular lifespan, we switched the cell growth medium from the standard 2% glucose condition to different glucose levels at the beginning of aging and tracked the lifespans of single cells (Fig. 2; Materials and Methods).

**Fig 2.**
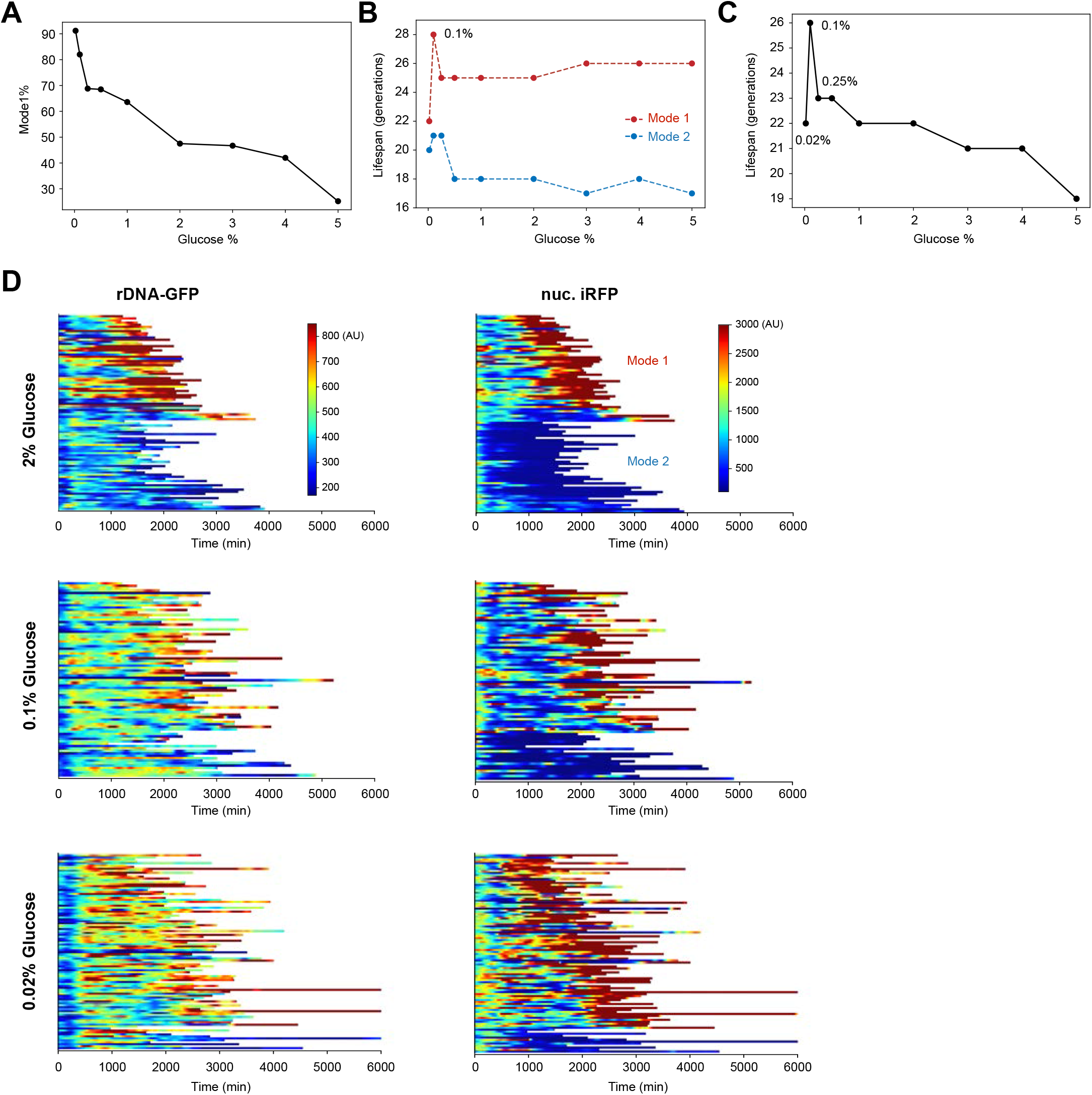
The effects of glucose levels on aging and lifespan. (**A**) The percentage of Mode 1 cells in an aging population as a function of glucose concentration. (**B**) The average lifespans of Mode 1 and Mode 2 cells as a function of glucose concentration (**C**) The average lifespan of a whole aging population as a function of the glucose level. Lifespan curves and statistical analysis are shown in Fig. S2. (**D**) Single-cell color map trajectories of rDNA-GFP (left) and nuclear-anchored iRFP (right) during aging under different glucose levels (2% Glucose: n = 91; 0.1% Glucose: n = 80; 0.02% Glucose: n = 100). Each row represents the time trace of a single cell throughout its life span. Color represents the fluorescence intensity as indicated in the color bar. Color maps for rDNA-GFP and nuc. iRFP are from the same cells with the same vertical order. For each glucose level, cells are classified into Mode 1 (top) and Mode 2 (bottom) based on their aging phenotypes and reporter dynamics.

We observed that changing glucose levels dramatically affected the fate decision in aging – decreased glucose promoted Mode 1 aging, whereas increased glucose promoted Mode 2 aging; the glucose level negatively correlated with the proportions of Mode 1 vs Mode 2 in an aging population (Fig. 2A). However, the average lifespans within Mode 1 and Mode 2 aging cells remained relatively unchanged (Fig. 2B). The exception was with 0.1% glucose, which markedly extended the lifespans of both Mode 1 and Mode 2 aging cells. Further decreasing the glucose level to 0.02%, however, largely shortened the lifespan of Mode 1 cells (>90% of the population). As a result, 0.1% glucose appeared to be an optimal level of CR that maximally extended the lifespan in yeast (Fig. 2C; Lifespan curves and t-tests are included in Fig. S2).

To further define the lifespan-extending effect of the optimal CR conditions, we monitored rDNA silencing loss and heme depletion, two major age-induced deterioration processes that regulate lifespan (Li et al., 2020; Zhou et al., 2023). To track rDNA silencing, we used a GFP reporter inserted within the rDNA (rDNA-GFP) (Li et al., 2017). Because expression of the reporter is subject to silencing, increased reporter fluorescence indicates reduced silencing, whereas decreased fluorescence indicates enhanced silencing. To track heme biogenesis, we used a nuclear-anchored iRFP reporter (nuc. iRFP) (Filonov et al., 2011). Its fluorescence depends on a heme degradation product, biliverdin, and thereby correlates with cellular heme levels (Li et al., 2020). We observed that, under the standard 2% glucose condition, about half of cells aged in Mode 1 with decreased rDNA silencing (as indicated by increased rDNA-GFP signal) and increased heme biogenesis (as indicated by increased iRFP signal) at late stages of aging, whereas the other half aged in Mode 2, with decreased heme abundance and modestly changed rDNA silencing (Fig. 2D, top), consistent with previous reports (Li et al., 2020; Zhou et al., 2023).

In contrast, in 0.1% glucose, the majority (82%) of cells exhibited Mode 1 aging, but these cells showed a substantial time delay in the loss of rDNA silencing and elevation of heme biogenesis (Fig. 2D, middle), compared to Mode 1 cells in 2% glucose. Decreasing the glucose level to 0.02% further increased the proportion of Mode 1 aging cells (91%). However, we observed an earlier occurrence of age-induced rDNA silencing loss and heme elevation (Fig. 2D, bottom), compared to Mode 1 cells in 0.1% glucose, in accord with a shorter lifespan under this condition. These results suggest that the maximal lifespan extension under 0.1% glucose could be attributed, at least partially, to the time delay in rDNA silencing loss and heme elevation during aging.

### The emergence of a “longevity fixed point” by CR

To quantitatively analyze mechanisms underlying the emergence of an optimal CR condition, we expanded a recently proposed model, comprising the mutual inhibition circuit of Sir2 and HAP that resembles a toggle switch to drive the fate decision in yeast aging (Li et al., 2020). Previous studies revealed that CR can elevate the levels of both Sir2 and HAP in yeast (Anderson et al., 2003; DeRisi et al., 1997; Lin et al., 2002; Medvedik et al., 2007; Westholm et al., 2008). To incorporate these effects in the model, the total amounts of Sir2 and HAP were considered as the functions of the glucose level (Fig. 3, A and B). Thus,

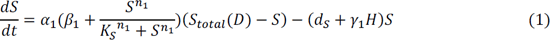

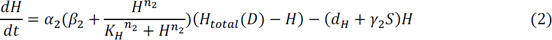

**Fig. 3.**
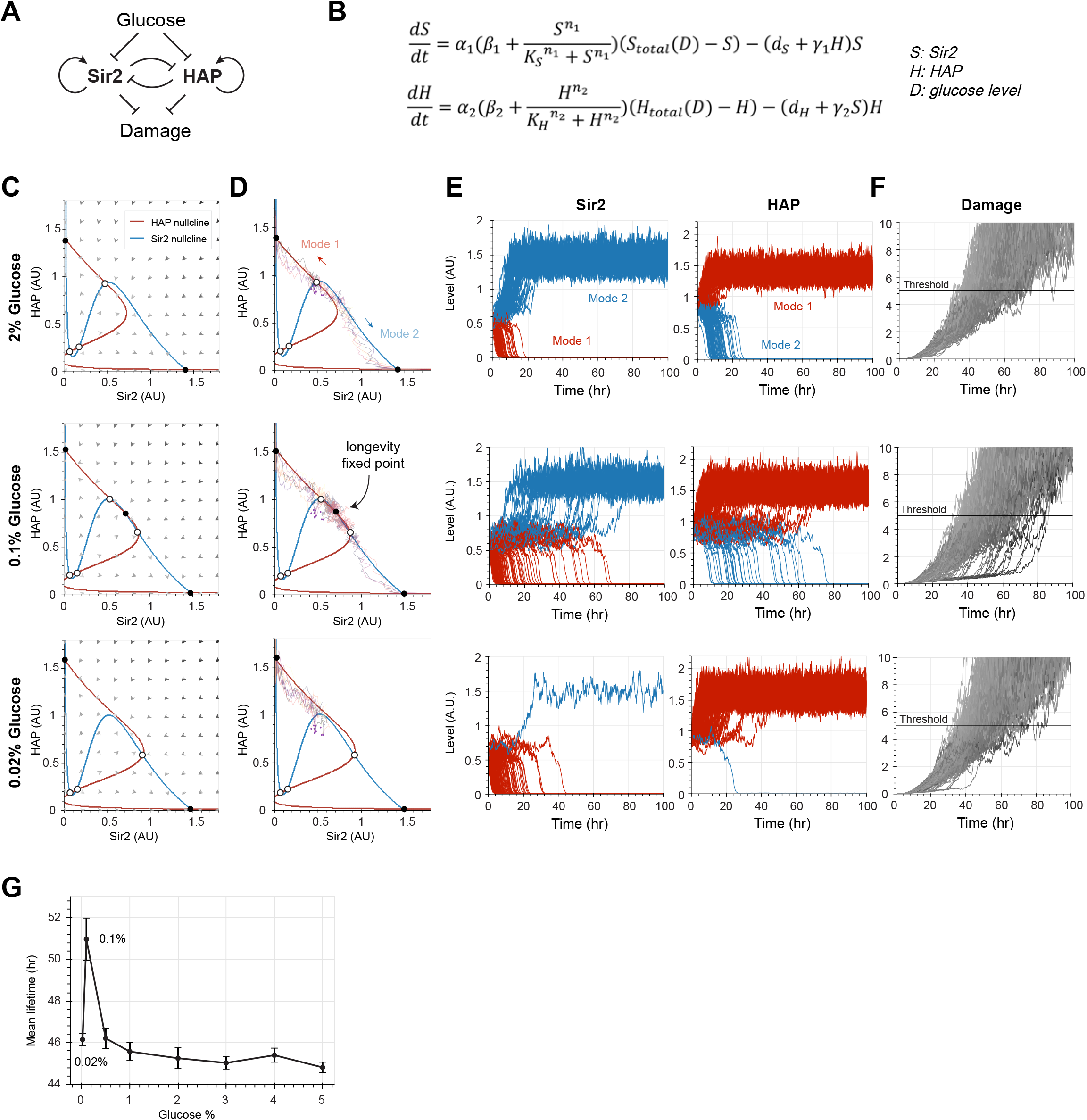
Computational modeling unravels the stability changes of the Sir2-HAP circuit under different glucose levels. (**A**) Diagram of the circuit topology. (**B**) Equations of the model. *S* is the concentration of enzymatically active Sir2, *H* is the concentration of active HAP complex and *D* is the concentration of glucose. *S_total_* and *H_total_* are piecewise functions of *D*, representing the total concentration of Sir2 and HAP, respectively (see Methods for details). (**C**) Phase planes demonstrate the stability changes of the Sir2-HAP circuit under different glucose levels. The nullclines of Sir2 and HAP are represented in blue and red, respectively. The arrows represent the rate and direction of the system’s movement. Fixed points are indicated with open (unstable) and closed (stable) circles. The stable fixed point at the top left corner of each phase plane corresponds to the terminal state of Mode 1 aging and the one at the bottom right corner corresponds to the terminal state of Mode 2 aging. At 0.1% glucose, a third stable fixed point (“longevity fixed point”) emerges with intermediate levels of Sir2 and HAP. (**D**) Stochastic simulations of aging trajectories on the Sir2-HAP phase planes under different glucose levels. The initial points of each trajectory (purple, n=10) were randomly generated based on the Gaussian distribution for simulation. Trajectories were numerically calculated by the stochastic version of equations from (B). (**E**) Stochastic simulations of the Sir2 and HAP time traces during aging under different glucose levels. The initial points (n=200) were randomly generated based on the Gaussian distribution for simulation. Red and blue curves represent the time traces of Mode 1 and Mode 2 cells, respectively. (**F**) Simulated time traces of damage accumulation based on Sir2 and HAP dynamics in (E). The cell is considered “dead” once its damage level exceeds the arbitrary threshold (horizontal line). (**G**) Simulated lifetime as a function of the glucose concentration. Simulations were performed 5 times with 100 cells for each simulation. The black dots and error bars represent the mean value and standard deviation of simulated lifetimes, respectively.

Here, *S*, *H* are concentrations of Sir2 and HAP, respectively; *α*_1_, *α*_2_ are maximum production rates for the positive feedback loop of Sir2 and HAP; *β*_1_, *β*_2_ are basal activation factors of Sir2 and HAP; *K*_S_, *K*_H_ are half-activation constants of Sir2 and HAP; *d*_*S*_, *d*_*H*_ are degradation rates of Sir2 and HAP; *γ*_1_, *γ*_2_ are repressive strength of HAP and Sir2; *n*_1_, *n*_2_ are Hill coefficients for Sir2 and HAP autoregulation. Total capacity of Sir2 and HAP are defined as piecewise functions of glucose concentration *D*:

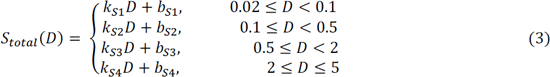

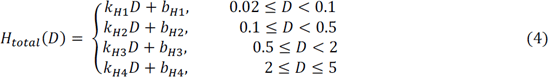

The parameters of the model were estimated by fitting the simulations with the data on the ratios of Mode 1 vs Mode 2 under different glucose conditions (Table S4).

To study the stochastic dynamics of aging, we used the following Langevin equations by adding noise terms to the deterministic equations:

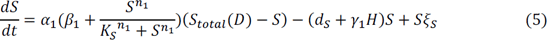

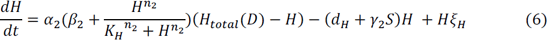

The noise terms *ξ*_*S*_, *ξ*_*H*_ are uncorrelated white Gaussian processes with zero mean and autocorrelation 〈*ξ*_*i*_(*t*)*ξ*_*i*_(*t*′)〉 = *ε*_*i*_*δ*(*t* − *t*′); *i* ∈ {*H*, *S*}, where *δ*(*t* − *t*′) is Dirac’s delta function and *ε*_*i*_ is the magnitude of *i*.

The effects of glucose levels on the dynamics of Sir2 and HAP can be analyzed graphically by plotting the nullclines and vector field in a Sir2-HAP phase plane under different concentrations of glucose (Fig. 3, C and D; Movie S1). With the standard 2% glucose level, the system has two stable fixed points – the low Sir2, high HAP point that corresponds to the terminal state of Mode 1 aging and the high Sir2, low HAP point that corresponds to the terminal state of Mode 2 aging (Fig. 3, C and D, top). Stochastic simulations showed that roughly equal numbers of cells progressed toward either of these two steady states during aging (Fig. 3E, top). When the glucose concentration decreases, an unstable fixed point between the two stable fixed points moves toward the high Sir2, low HAP region, biasing the fate decision toward the low Sir2, high HAP state and thereby Mode 1 aging (Movie S1). A third stable fixed point emerges in the intermediate Sir2, intermediate HAP region of the phase plane when the glucose concentration reaches a certain level (Fig. 3, C and D, middle). Stochastic simulations showed that a large fraction of cells fluctuated around this new stable fixed point for a period of time before eventually deviating to the low Sir2, high HAP or high Sir2, low HAP states, driven by the noise (Fig. 3E, middle). These fluctuations are in accord with the delayed rDNA silencing loss and heme elevation observed experimentally under the optimal CR condition (0.1% glucose) (Fig. 2D, middle). This new stable fixed point requires a subtle balance between Sir2 and HAP and hence disappears quickly when the glucose concentration is further decreased (Fig. 3, C and D, bottom). As a result, at 0.02% glucose, the majority of cells aged with Mode 1 aging and rapidly approached the low Sir2, high HAP state (Fig. 3E, bottom), consistent with the aging dynamics observed under this condition (Fig. 2D, bottom).

To simulate the effects of glucose alterations on lifespan, we linked the Sir2-HAP circuit to cell death using a paradigmatic model framework recently developed by Uri Alon’s group (Karin et al., 2019; Yang et al., 2023), in which the aging process is described as a competition between accelerating damage accumulation and saturating damage removal. The cell death occurs when the level of intracellular damage exceeds a certain threshold value. In our model, we assume that the damage removal rate is dependent on the levels of Sir2 and Hap4. Specifically, the equation for the damage *ψ* is written as follows:

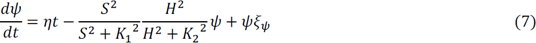

Here, *η* is the production rate of damage and *ξ*_*ψ*_ is Gaussian white noise term with strength *ε*_*ψ*_. Initial conditions for Sir2 and HAP follow a Gaussian probability distribution 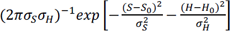 (see Table S4 for the parameters of simulations).

Decreasing either Sir2 or HAP levels reduces damage repair/removal and thereby accelerates damage accumulation. The lifespan of a cell is calculated as the time taken for its damage level to hit the threshold (Fig. 3F). The optimal CR condition allows for the system to spend an extended period near the new stable fixed point with intermediate Sir2 and HAP levels and hence to slow damage accumulation, leading to the maximally extended lifespan across different glucose levels (Fig. 3G). We therefore designated this stable fixed point as a “longevity fixed point.”

### Sir2 overexpression can stabilize the longevity fixed point

To determine the dependence of the longevity fixed point on the glucose concentration, we performed a bifurcation analysis of our model (Equations 5 and 6) and showed that, in the WT system, the longevity fixed point emerges within a narrow range of glucose levels, into which the optimal CR falls (Fig. 4A, left). Consistently, in WT, the longevity fixed point only exists within a small triangular region of the Sir2-HAP parameter space and the trajectory depicting glucose level changes passes across only the tip of that region (Fig. 4A, right).

**Fig. 4.**
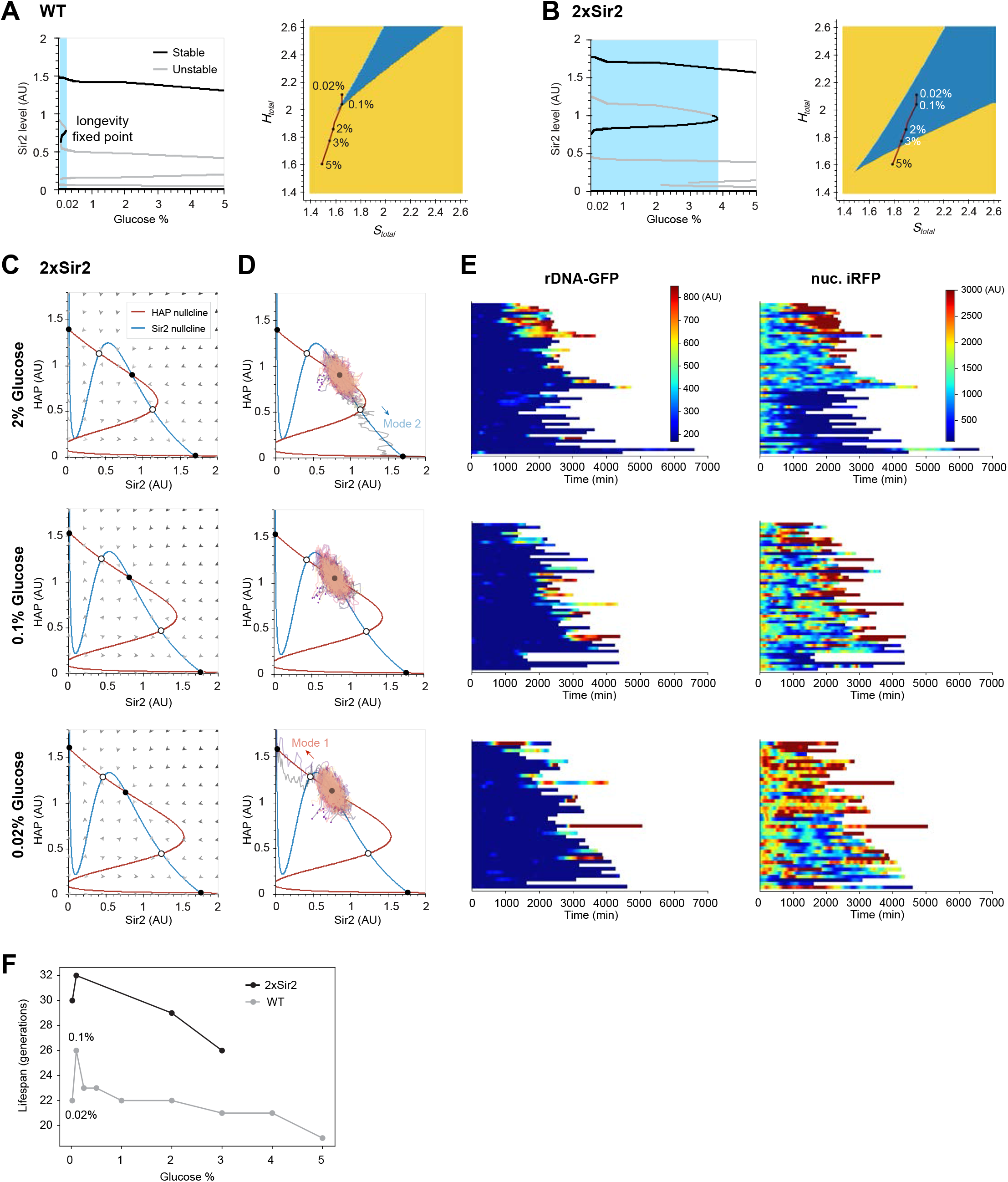
Sir2 overexpression stabilizes the longevity fixed point. (**A-B**) Dependence of the longevity fixed point on the glucose level in WT and 2xSir2. (A-B, left) Bifurcation analysis for the stability of the Sir2-HAP circuit as a function of the glucose level. Black lines and grey lines represent stable and unstable fixed points, respectively. The range of glucose levels in which the longevity fixed point exists is shaded in blue. (A-B, right) The dependence of the longevity fixed point on the total amounts of Sir2 (*S_total_*) and HAP (*H_total_*). The glucose level can affect the values of *S_total_* and *H_total_*, which is depicted by the red line. The blue region represents the area where the longevity fixed point exists in the parameter space. (**C**) The Sir2-HAP phase planes for 2xSir2 under different glucose levels. (**D**) Stochastic simulations of aging trajectories on the Sir2-HAP phase planes for 2xSir2 under different glucose levels. (**E**) Single-cell color map trajectories of rDNA-GFP (left) and nuc. iRFP (right) for cells with 2-fold overexpression of *SIR2* under different glucose conditions (2% glucose: n = 53; 0.1% glucose: n = 50; 0.02% glucose: n = 41). (**F**) The experimentally-measured lifespans of WT vs 2xSir2 as a function of the glucose level.

Our previous analysis of the Sir2-HAP model revealed that 2-fold overexpression of Sir2 can also create the longevity fixed point under standard growth conditions (corresponding to long-lived Mode 3 aging) (Li et al., 2020). We therefore examined how Sir2 overexpression can alter the dependence of this fixed point on glucose levels. Our model predicted that increasing Sir2 abundance by 2-fold (see Materials and Methods) can reshape the phase diagram of the system so that the longevity fixed point can exist within a much wider range of glucose levels and a larger region of the Sir2-HAP parameter space (Fig. 4B).

In addition, because the two flanking unstable fixed points are moved further away from the longevity fixed point by Sir2 overexpression, the system becomes less likely to deviate to the other two “aged” stable fixed points by the noise (Fig. 4, C and D), which could lead to longer lifespans under a wide range of glucose levels, compared to that of WT cells under the optimal CR condition. Changing the glucose level can still modulate the distances between the longevity fixed point and the two flanking unstable fixed points and thereby can influence the fate decision in the 2 x Sir2 system: 2% glucose biased the system toward deviation to the high Sir2, low HAP state and Mode 2 aging, whereas 0.02% glucose biased the system toward the low Sir2, high HAP state and Mode 1 aging; 0.1% glucose remained the optimally balanced condition with the least chance of deviations.

To test these predictions experimentally, we overexpressed Sir2 by 2-fold and monitored cell aging under different glucose concentrations. Consistent with the model, we observed substantially delayed rDNA silencing loss and heme elevation in a large fraction of cells under all the tested glucose concentrations (0.02% - 3%) (Fig. 4E), which resulted in longer lifespans than that of WT cells over a wide range of glucose levels (Fig. 4F; lifespan curves and t-tests are included in Fig. S3). These results are in accord with a more stable longevity fixed point in the 2 x Sir2 system. In addition, glucose levels can indeed affect the fate decisions of aging cells with Sir2 overexpression (Fig. 4E), consistent with model simulations (Fig. 4D).

### External glucose oscillations can enable “dynamic stabilization” of the aging process

We recently engineered a synthetic Sir2-HAP gene oscillator that could slow cell deterioration, resulting in a dramatically extended lifespan (Zhou et al., 2023). In this study we considered the possibility that oscillatory glucose inputs may be able to drive dynamic stabilization of the Sir2-HAP circuit and thereby promote longevity. For example, in our model, the system does not have the longevity fixed point at either 2% or 0.02% glucose levels and is unstable in the intermediate Sir2, intermediate HAP range. However, oscillations between 0.02% and 2% glucose levels may be able to restrict the system to that region, reaching a state of dynamic stabilization.

To test this possibility *in silico*, we first performed stochastic simulations with external glucose oscillations between 0.02% and 2%. We found that at least a fraction of cells indeed remains within the intermediate Sir2, intermediate HAP state for an extended period before escaping to the low Sir2 or low HAP states (Fig. 5, A-C; Movie S2). Increasing the frequency of glucose oscillations increases the proportion of cells that exhibit such a delay (Fig. 5D; Fig. S4).

**Fig. 5.**
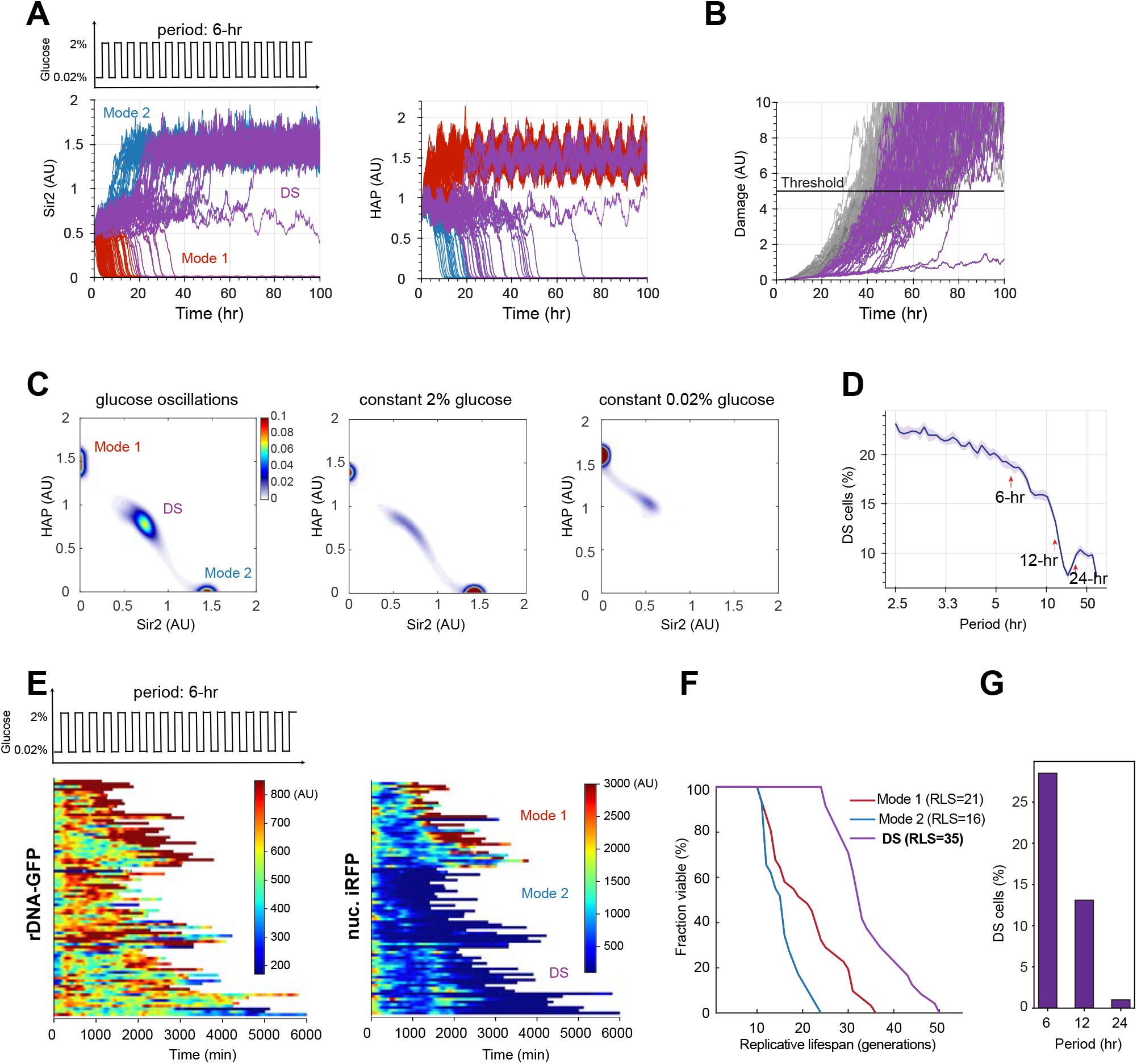
External glucose oscillations enable dynamic stabilization of the aging process. (**A**) Stochastic simulations of the Sir2 and HAP time traces during aging under glucose oscillations between 0.02% and 2%, with a 6-hr period (n=200). Dynamically stabilized (DS) cells were defined as those that continue to fluctuate around the intermediate Sir2 and HAP state for an extended period before deviation to Mode 1 or Mode 2 (see Material and Methods). The time traces of DS cells are shown in purple. (**B**) Simulated time traces of damage accumulation based on Sir2 and HAP dynamics in (A). The cell is considered “dead” when its damage level exceeds the arbitrary threshold (horizontal line). (**C**) Stability of aging trajectories upon glucose oscillations vs constant glucose levels. Stochastic simulations were performed in a Sir2-HAP plane of 200×200 grids, and the average retention probability in each grid was computed, as indicated by the color bar. (**D**) The percentage of DS cells in an aging population as a function of the period of glucose oscillations, from model simulations. (**E**) Single-cell color map trajectories of rDNA-GFP (left) and nuc. iRFP (right) during aging under glucose oscillations between 0.02% and 2% with a period of 6 hours (n=84). Each row represents the time trace of a single cell throughout its life span. Color represents the fluorescence intensity as indicated in the color bar. Color maps for rDNA-GFP and nuc. iRFP are from the same cells with the same vertical order. Under each glucose level, cells are classified into Mode 1 (top), Mode 2 (middle) and DS (bottom) based on their aging phenotypes and reporter dynamics. (**F**) Lifespan curves of Mode 1, Mode 2, and DS cells. (**G**) The percentage of DS cells in an aging population as a function of the period of glucose oscillations, determined by experiments.

To determine whether this delay depends on the existence of the stable longevity fixed point at the constant 0.1% glucose condition, we modified the dependence functions of Sir2 and HAP on the glucose level, *S*_*total*_(*D*) and *H*_*total*_(*D*), so that the longevity fixed point would never emerge in the system at any glucose level (Fig. S5A). We found that external glucose oscillations can still generate an extended delay before fate commitments (Fig. S5B), indicating that the dynamic stabilization does not require the presence of the stable longevity fixed point at an intermediate level of glucose. The mechanistic reason for the emergence of the delay is that oscillations between two different levels of glucose lead to an averaging effect of the two corresponding states in the (*S*_*total*_, *H*_*total*_) parameter space and this novel averaged state features a stable fixed point for longevity (Fig. S5A). These modeling results suggest that oscillatory external inputs can enable dynamic stabilization of Sir2 and HAP levels within an optimal range and thereby extend lifespan.

To test these computational predictions experimentally, we integrated our microfluidic device with a computer-controlled electrovalve (Hansen et al., 2015; Hersen et al., 2008) to deliver dynamic patterns of environmental inputs into the culture chambers. We applied external glucose oscillations between 0.02% and 2% with 6-hour period throughout the lifespans of yeast cells and tracked their aging processes. In addition to Mode 1 and Mode 2 aging cells, we observed that about one-third of cells showed fluctuating rDNA-GFP and iRFP signals for a long period of time before deviating to the low HAP state (Fig. 5E). These cells corresponded to the fraction of cells with dynamic stability in our model simulations. We hence designated them as dynamically stabilized (DS) cells. These DS cells turned out to be very long-lived, with an average lifespan of 35 generations, much longer than that of Mode 1 or Mode 2 cells from the same aging population (Fig. 5F). We further tested glucose oscillations with an input period of 12 hours and 24 hours. Consistent with model simulations, decreasing the input frequency reduced the proportion of Mode DS cells in an aging population (Fig. 5G).

Taken together, these results validated our modeling results and demonstrated that external glucose oscillations can indeed enable dynamic stabilization of aging processes, leading to a prolonged lifespan.

## Discussion

In this study, we devised a simple mathematical model that can not only reproduce the single-cell aging data under variable glucose conditions but also shed light on the underlying biological mechanisms governing the metabolic regulation of aging. Importantly, the model has predictive power that can be used to guide the design of interventional strategies for longevity and to reveal the theoretical principles underlying these strategies. In particular, we focused on the Sir2-HAP toggle switch circuit that mediates the progression of two major aging paths in single yeast cells – one leads to a low Sir2, high HAP state, resulting in nucleolar decline, and the other leads to a high Sir2, low HAP state, causing mitochondrial deterioration (Fig. 3) (Li et al., 2020). Because both of these terminal states of aging are associated with markedly accelerated damage accumulation, the overall goal for pro-longevity interventions is to avoid, or at least delay, the fate commitment and progression toward these two detrimental steady states (stable fixed points) of the natural aging system. This is challenging in that interventions that elevate either end of the toggle, Sir2 or HAP, will simply push the cell to the other path toward aging and death.

Our model-based analysis here unraveled two general approaches to promote longevity: (1) create and stabilize a new longevity fixed point in the “healthy” intermediate Sir2 and HAP region; and (2) enable dynamic stabilization of the system around the “healthy” state. Both can be realized by dynamically modulating environmental glucose levels.

For the first approach, the longevity fixed point, which achieves a subtle balance between Sir2 and HAP at intermediate levels, can be stabilized by either an optimal level of CR or moderate overexpression of Sir2. Both interventions enable the cell to maintain a state with intermediate levels of Sir2 and HAP for an extended period of time. Among them, genetic manipulation of Sir2 leads to a fixed point that is less sensitive to environmental glucose alterations (Fig. 4B). Intriguingly, however, although Sir2 overexpression and CR extend the lifespan through the same mechanism, their effects can still be additive (Fig. 4F). The glucose level, by acting on HAP as well as Sir2, can modulate the stability of the longevity fixed point generated by Sir2 overexpression and thereby influence the lifespan (Fig. 4D). As an immediate application, our model can be used to design the combined treatments with CR and pharmacological activators of sirtuins, in which *in silico* simulations could help refine the optimal dosage combinations for maximizing the effects on lifespan extension.

For the second approach inspired by the nonlinear dynamics and control theory, we adopted the concept of dynamic stabilization for designing pro-longevity strategies. Indeed, an unstable state of a dynamical system, such as the natural aging circuit with intermediate levels of Sir2 and HAP, can be stabilized by applying periodic oscillations of external glucose levels at certain frequencies (Fig. 5, A and B). More generally, it is possible that oscillatory inputs can stabilize the system in a healthy, long-lived state whereas the constant inputs cannot (Fig. 5C). It is interesting to note that this method of dynamical stabilization is reminiscent of the stabilization of inverted pendulum by periodic vibrations of its pivot point, so–called Kapitsa’s pendulum (Blackburn, 1992). Tuning the frequency can modulate the strength of this dynamic stabilization and thereby the proportion of cells that age through this long-lived state of unstable equilibrium (Fig. 5D).

Temporally-varying metabolic interventions, such as intermittent fasting, are attracting increasing attention as a potential approach to effectively promote longevity and healthspan (Fontana and Partridge, 2015; Hwangbo et al., 2020; Longo and Panda, 2016; Mattson et al., 2017), but definition of mechanisms underlying their effectiveness remain elusive. Our dynamic stabilization theory of aging, validated here in yeast, may provide the theoretical underpinning for the effects of temporally-varying interventions. As large-scale intervention testing is prohibitively time- and resource-intensive, models based on this theoretical framework can help screen and identify *in silico* the optimal dynamic patterns of metabolic interventions for experimental examinations. Such model-directed approaches can also be applied to identify time-based administration regimes for a series of metabolic intervention compounds named “caloric restriction mimetics” (CRM), including rapamycin, metformin, and spermidine, shown to increase lifespan in multiple model organisms (Barzilai et al., 2016; Madeo et al., 2019; Madeo et al., 2018; Martel et al., 2021; Miller et al., 2011; Mohammed et al., 2021; Novelle et al., 2016; Selvarani et al., 2021).

In addition to the two approaches presented here, a third approach, creating a limit cycle for longevity, was conceived by attempts to modify the circuit structure in the model. Rewiring the mutual inhibition between Sir2 and HAP into a negative feedback loop can replace the detrimental stable fixed points with a limit cycle on the Sir2-HAP phase plane, leading to sustained oscillations in Sir2 and HAP levels and thereby avoiding fate commitment to either low Sir2 or low HAP state. This strategy has been realized by genetic circuit engineering in the cell (Zhou et al., 2023).

Our model is designed specifically for yeast aging, yet due to its abstract nature, it can be readily applied to design interventions for any fate decision processes driven by toggle switch circuits, such as trophectoderm differentiation by the Oct3/4-Cdx2 circuit (Niwa et al., 2005), induced pluripotent stem cell reprogramming by the Oct4-Sox2 circuit (Shu et al., 2013), hematopoietic stem cell differentiation by the GATA-1 and PU.1 circuit (Liew et al., 2006), and many others. Future studies will be focused on identifying major toggle switch circuits that drive aging in other organisms or in human cells, based on which we can apply our modeling framework to design and test universal interventional strategies for promoting longevity.

## Acknowledgements

We thank Dr. Lorraine Pillus for insightful discussions and suggestions on the manuscript. This work was supported by National Institutes of Health R01AG056440 (to N.H., J.H., L.S.T.), R01GM144595 (to N.H., J.H., L.S.T.), R01GM111458 (to N.H.) and R01AG068112 (to N.H.).

## Author Contributions

Conceptualization, Y.L., Z.Z., L.S.T., J.H., and N.H.; Methodology, Y.L., Z.Z., L.S.T., J.H., and N.H.; Investigation, Y.L., Z.Z., S.W., G.N., A.Z.; Formal Analysis, Y.L., Z.Z., S.W., L.S.T.; Writing - Original Draft, Y.L., Z.Z., and N.H.; Writing - Review & Editing, Y.L., Z.Z., S.W., G.N., A.Z., L.S.T., J.H., and N.H.; Resources, L.S.T., J.H., and N.H.; Supervision, N.H.; Funding Acquisition, L.S.T., J.H., and N.H.

## Materials and Methods

### Strain and plasmid construction

Standard protocols were used for molecular cloning. All yeast strain used in this study were constructed based on BY4741 (*MAT*a *his3Δ1 leu2Δ0 met15Δ0 ura3Δ0*). Details of strains, plasmids, and primers are included in Table S1-S3.

To make the *SIR2* 2-fold overexpression plasmid, a XbaI_pSIR2_SIR2_EcoRI fragment containing 620 bp of the *SIR2* promoter + the *SIR2* ORF was made by PCR and then ligated into pRS303, yielding plasmid NHB0638.

To make the plasmid for the ATP reporter, a DNA fragment containing sequences from iATPSnFR^1.0^ (including epsilon subunit, Linkers 1 and 2, GFP sequences) (Lobas et al., 2019) was synthesized by IDT with codon optimization for yeast BY4741. Then PCR and Gibson Assembly were used to insert the 408bp *TEF1* promoter and the iATPSnFR^1.0^ fragment into pRS303 plasmid, yielding NHB1222.

To make the plasmid for glycolysis (FBP) reporter, pHO_pTEFmut7_CggR_R250A_ble (#124585) and pCggRO-reporter (#124582) plasmids were purchased from Addgene (Monteiro et al., 2019). Then forward primer with BamHI cutting site and reverse primer with EcoRI cutting site were used to PCR CggRO sequence. Forward primer with XhoI cutting site and reverse primer with EcoRI cutting site were used to PCR pTEFmut7_CggR_R250A sequences. After that, the two PCR fragments were ligated into pRS303 plasmid cut by XhoI and BamHI to get the plasmid NHB1227.

The yeast strain with the nuc. iRFP reporter (NH268) and the strain with both nuc. iRFP and the NTS1 silencing reporters (NH270) was made as previously described (Li et al., 2017). The yeast strain with the ATP reporter was made by transforming NH268 with DNA fragments from NHB1222 digested by BsmI. The yeast strain with Hxt1-CFP, Hxt2-YFP, Hxt3-mCherry reporters was made by the following steps: firstly, mCherry-LEU2 was amplified by PCR and integrated at the C-terminus of Hxt3 by homologous recombination to make the Hxt3-mCherry strain; secondly, CFP-HIS3 was amplified by PCR and integrated at the C-terminus of Hxt1 by homologous recombination to make the Hxt3-mCherry, Hxt1-CFP strain; Lastly, YFP-URA3 was amplified by PCR and integrated at the C-terminus of Hxt2 by homologous recombination to make the Hxt3-mCherry, Hxt1-CFP, Hxt2-YFP strain. Similarly, the yeast strain with the Hxk1-GFP reporter was made by transforming GFP-HIS3 fragments amplified from PCR. The yeast strain with the glycolysis reporter was made by transforming NH268 with DNA fragments from NHB1227 digested by BsmI.

### Setting up microfluidic experiments and time-lapse microscopy

The microfluidic devices and experiments were set up as previously described (Zhou et al., 2023). The yeast cells were incubated in SC medium containing 2% glucose before loading into the microfluidic devices. SC media containing different concentrations of glucose were delivered to cells via the media supply syringes after cell loading and were applied throughout the experiments.

A computer-controlled electrovalve was used to deliver external glucose oscillations to aging cells in the microfluidic device. The 3-way 0.054 ports electrovalve (The Lee Company, #LFYA1226032H) was connected to a 4-channel USB powered relay module (Numato Lab, SKU: USBPOWRL004). Each channel of the relay could be programmed to control the media input to one microfluidic device by custom-designed MATLAB App. To generate glucose oscillations, growth media with 2% glucose and 0.02% glucose were connected to Port I and O of the electrovalve, respectively, which can be computer-controlled. The outlet port of the valve was connected to the inlet of the microfluidic device.

Time-lapse microscopy experiments were conducted using a Nikon Ti-E inverted fluorescence microscope with an EMCCD camera (Andor iXon X3 DU897). The light source is a spectra X LED system. Images were taken using a CFI plan Apochromat Lambda DM 60X oil immersion objective (NA 1.40 WD 0.13MM). In all experiments, the images were acquired for each fluorescence channel every 15 min for a total of 90 to 120 hours. The exposure and intensity setting for each channel were set as follows: 1) For the glucose tuning assay: Phase 80 ms, GFP 4 ms/30ms at 10% lamp intensity with an EM Gain of 70, and iRFP 50 ms at 15% lamp intensity with an EM Gain of 300; 2) For detecting the ATP reporter : Phase 80 ms, GFP 30ms at 10% lamp intensity with an EM Gain of 70, and iRFP 300ms at 15% lamp intensity with an EM Gain of 300; 3) For detecting Hxt1-3: Phase 80 ms, mCherry 100ms at 25% lamp intensity with an EM Gain of 250, YFP 100ms at 10% lamp intensity with an EM Gain of 250, CFP 60ms at 10% lamp intensity with an EM Gain of 250 and iRFP 300ms at 15% lamp intensity with an EM Gain of 100; 4) For detecting the FBP reporter : Phase 50 ms, YFP 200ms at 10% lamp intensity with an EM Gain of 250, and iRFP 300ms at 15% lamp intensity with an EM Gain of 100.

### Quantification of single-cell aging traces

Image processing was conducted using a custom MATLAB code (Li et al., 2020; Li et al., 2017; Zhou et al., 2023). The background of images from each fluorescence channel were subtracted. Cell nuclei were masked by thresholding iRFP signal. The mean intensity value of the top 40% of the pixels of fluorescence reporters was quantified, as described previously (Li et al., 2020; Li et al., 2017; Zhou et al., 2023). The time traces of reporters were smoothed using the MATLAB function *smoothdata* with specification of the Gaussian method through a 15-element sliding window.

To plot the cell cycle length changes as a function of the percentage of lifetime, the vector of cell cycle length was interpolated to a new vector of 100 elements at evenly distributed 100 query points. For single cell aging dynamics and replicative lifespan (RLS) analyses, we collected data from at least 3 independent experiments. Any cells showing obvious abnormal morphologies upon cell loading were filtered out for RLS analysis. Any cells showing dislocation of reporter mask were filtered out for time trace analysis but included in RLS analysis. Significant numbers for RLS changes were calculated with Gehan-Breslow-Wilcoxon test by using Prism GraphPad 7 (GraphPad Software, CA).

### Computational Modeling

#### Stochastic simulations

The stochastic dynamics of aging were studied using the following Langevin equations by adding noise terms to the deterministic equations:

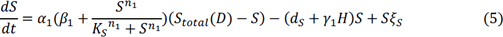

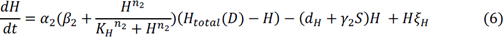

The noise terms *ξ*_*S*_, *ξ*_*H*_ are uncorrelated white Gaussian processes with zero mean and autocorrelation 〈*ξ*_*i*_(*t*)*ξ*_*i*_(*t*′)〉 = *ε*_*i*_*δ*(*t*− *t*′); *i* ∈ {*H*, *S*}, where *δ*(*t* − *t*′) is Dirac’s delta and *ε*_*i*_is the magnitude of *i*.

To simulate the lifetime of each cell, we coupled the saturated-repair stochastic equation described by (Yang et al., 2023) with equations (5) & (6). The damage removal term was modified with respect to the level of Sir2 and HAP, the equation is listed as following:

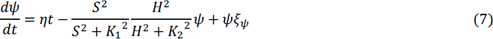

where *η* is the maximum production rate of damage, *ξ*_*ψ*_ is a Gaussian white noise term with strength *ε*_*ψ*_. The cell was defined as dead when cellular damage *ψ* reached the threshold which was set to 5 in our cases. Initial conditions for Sir2 and HAP follow a Gaussian probability distribution 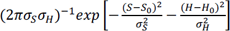 (see Table S4 for the parameters of simulations).

#### Bifurcation analysis

The fixed points of equation (1) & (2) were calculated by *fsolve* from SciPy. Points with negative values were omitted. The eigenvalues of Jacobian matrix of equations (1) & (2) at fixed point were calculated to determine its stability. The sweet spot (the third stable fixed point) was defined as the point that has both negative eigenvalues and lies within the range of 15%-70% of Sir2 and HAP total amount.

#### Mode ratio determination

Mode 2 was defined as *H* equals 0 at the end of simulated lifetime. We defined the number of Mode 1 cell equals *N-M_2_*, where the *N* is the number of overall cells and *M_2_* is the number of Mode 2 cells. The mode ratio equals 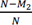.

#### Mode DS determination

To get the distributions of lifetime of Mode 1 and Mode 2 cells, 20,000 rounds of simulation were conducted corresponding to the conditions of 0.02% and 2% glucose, respectively. The lifetimes for Mode 1 and Mode 2 were fitted to gamma distributions separately (Fig. S4). The threshold in each mode for defining “Mode DS” was set as Cumulative Distribution Function (CDF)=95%.

#### 2 × Sir2 overexpression approximation

When another copy of *SIR2* gene was added, total Sir2 capacity was increased by a factor of κ. Equation 5 became:

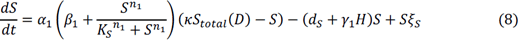

Considering Sir2 is nicotinamide adenine dinucleotide (NAD^+^)-dependent enzyme, overexpression of Sir2 may lead to competitive binding of NAD^+^, which reduces the inhibition strength (*γ*_2_) to HAP. The assumption is in line with our experimental results in which 2 × Sir2 in 2% glucose condition exhibited no effect on pushing the cells to Mode 2 (∼57% mode 1 of 2 × Sir2 vs ∼50% mode 1 of WT), indicating a compromise of increase of Sir2 capacity and reduction of Sir2 activity simultaneously. In addition, an extreme overexpression of Sir2 driven by the strong promoter, p*TDH3*, leads to almost 100% mode 1 cells. This suggests that Sir2 does not exert an inhibitory effect on HAP through the loss of Sir2 enzymatic activity, caused by severe competition for NAD^+^. In this case, we set κ equals 1.2 and *γ*_2_ equals 0.5.

---

**Table S1.** Strains used or constructed in this study. All strains will be available upon request.

| Strain Name | Description |
| --- | --- |
| NH0268 | *BY4741 MATa his3Δ1 leu2Δ0 met15Δ0 ura3Δ0, NHP6a-iRFP-kanMX* |
| NH0270 | *BY4741 MATa his3Δ1 leu2Δ0 met15Δ0 ura3Δ0, RDN1::NTS1-P<sub>TDH3</sub>-GFP-URA3,NHP6a-iRFP-kanMX* |
| NH0886 | *BY4741 MATa his3Δ1 leu2Δ0 met15Δ0 ura3Δ0, RDN1::NTS1-P<sub>TDH3</sub>-GFP-URA3,NHP6a-iRFP-kanMX, pSIR2::pSIR2-SIR2-HIS3* |
| NH1641 | *BY4741 MATa his3Δ1 leu2Δ0 met15Δ0 ura3Δ0, NHP6a-iRFP-kanMX, HXT3-mCherry::LEU2; HXT1-CFP::HIS3; HXT2-YFP::URA3* |
| NH1692 | *BY4741 MATa his3Δ1 leu2Δ0 met15Δ0 ura3Δ0, NHP6a-iRFP-kanMX, pTEF1-iATPSnFR1.0::HIS3* |
| NH1704 | *BY4741 MATa his3Δ1 leu2Δ0 met15Δ0 ura3Δ0, NHP6a-iRFP-kanMX, pTEFmut7-CggR_R250A-CggRO-YFP::HIS3* |
| NH1843 | *BY4741 MATa his3Δ1 leu2Δ0 met15Δ0 ura3Δ0, NHP6a-iRFP-kanMX, HXK1-GFP::HIS3* |

**Table S2.**
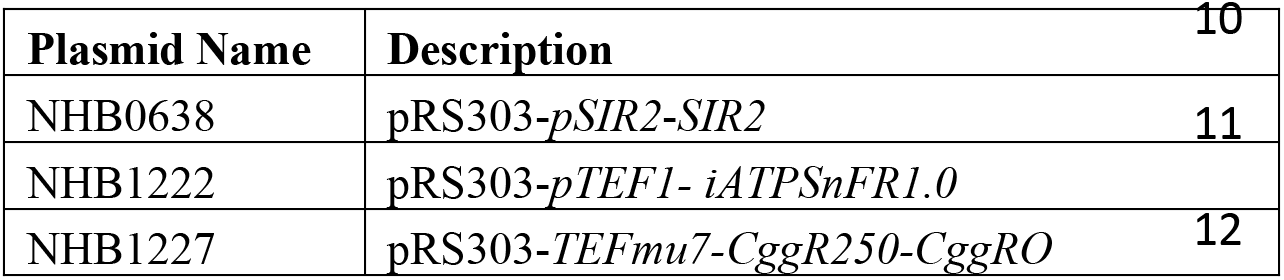
Plasmids constructed in this study. All plasmids will be available upon request.

**Table S3.**
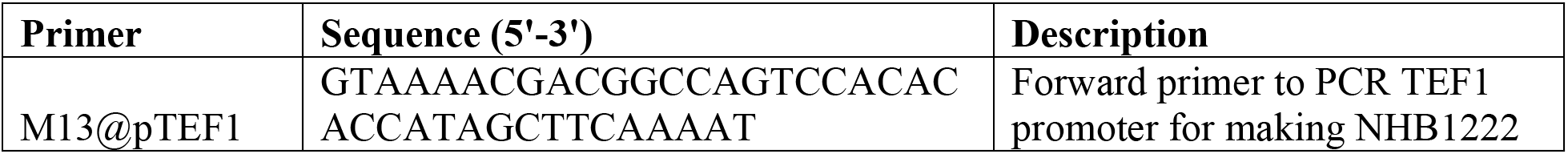
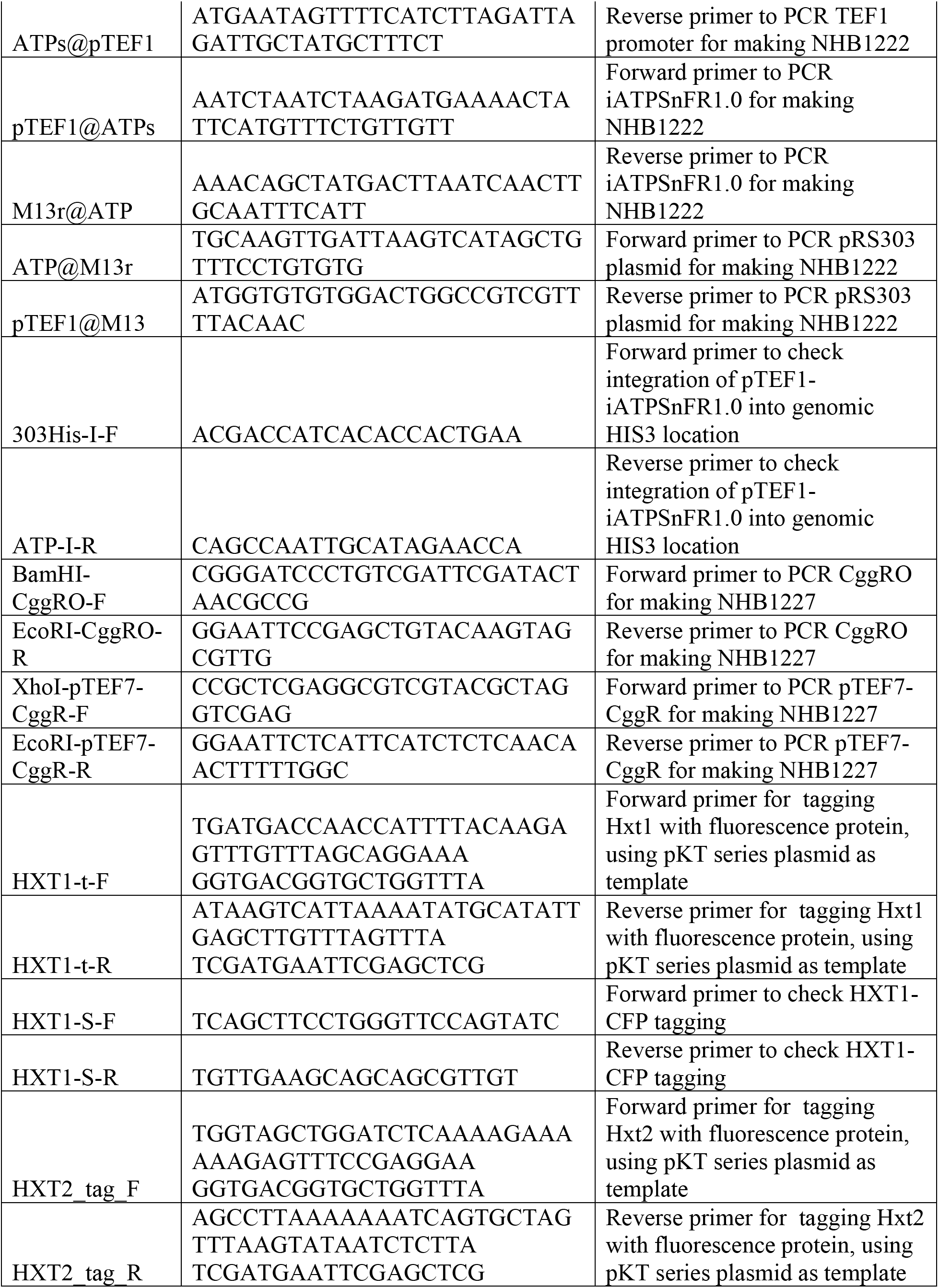
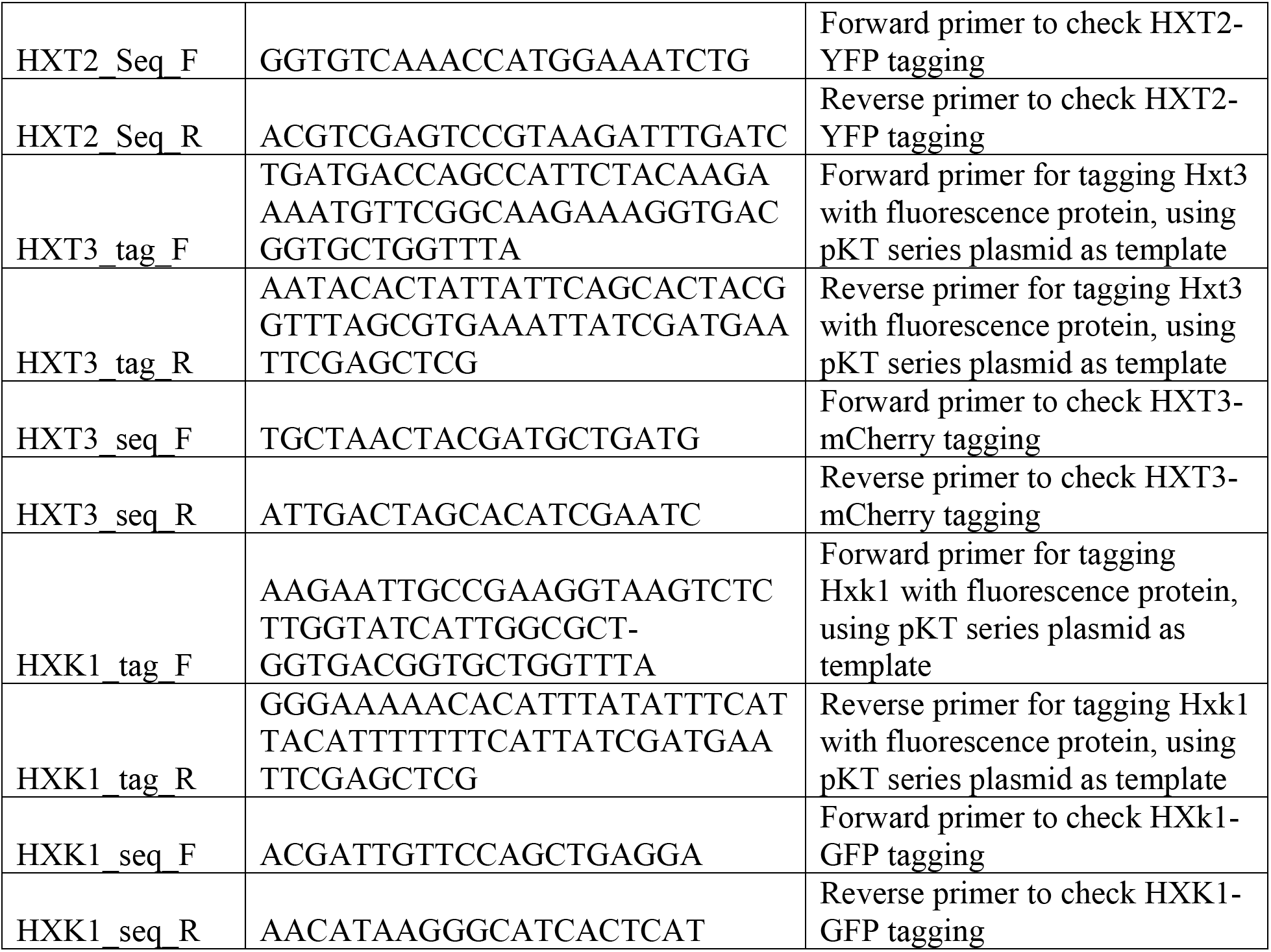
Primers used in this study.

**Table S4.** Parameters used in both deterministic and stochastic models.

| Parameter | Value |
| --- | --- |
| $\alpha_1$ | $1 \text{ h}^{-1}$ |
| $\alpha_2$ | $1 \text{ h}^{-1}$ |
| $\beta_1$ | $0.01 \text{ h}^{-1}$ |
| $\beta_2$ | $0.01 \text{ h}^{-1}$ |
| $K_S$ | 0.52 |
| $K_H$ | 0.62 |
| $d_S$ | $0.1 \text{ h}^{-1}$ |
| $d_H$ | $0.3 \text{ h}^{-1}$ |
| $\gamma_1$ | 1 |
| $\gamma_2$ | 1 |
| $n_1$ | 3 |
| $n_2$ | 3 |
| $k_{S1}$ | 0 |
| $k_{S2}$ | -0.15 |
| $k_{S3}$ | -0.007 |
| $k_{S4}$ | -0.03 |
| $b_{S1}$ | 1.65 |
| $b_{S2}$ | 1.66 |
| $b_{S3}$ | 1.59 |
| $b_{S4}$ | 1.64 |
| $k_{H1}$ | -0.87 |
| $k_{H2}$ | -0.32 |
| $k_{H3}$ | -0.03 |
| $k_{H4}$ | -0.085 |
| $b_{H1}$ | 2.13 |
| $b_{H2}$ | 2.07 |
| $b_{H3}$ | 1.93 |
| $b_{H4}$ | 2.03 |
| $\epsilon_H$ | 0.32 |
| $\epsilon_S$ | 0.32 |
| $\epsilon_\psi$ | 0.35 |
| $\eta$ | 0.005 |
| $S_0$ | 0.49 |
| $H_0$ | 0.86 |
| $\sigma_S$ | 0.05 |
| $\sigma_H$ | 0.05 |

