## Supplemental figures for "Enhanced cellular longevity arising from environmental fluctuations"

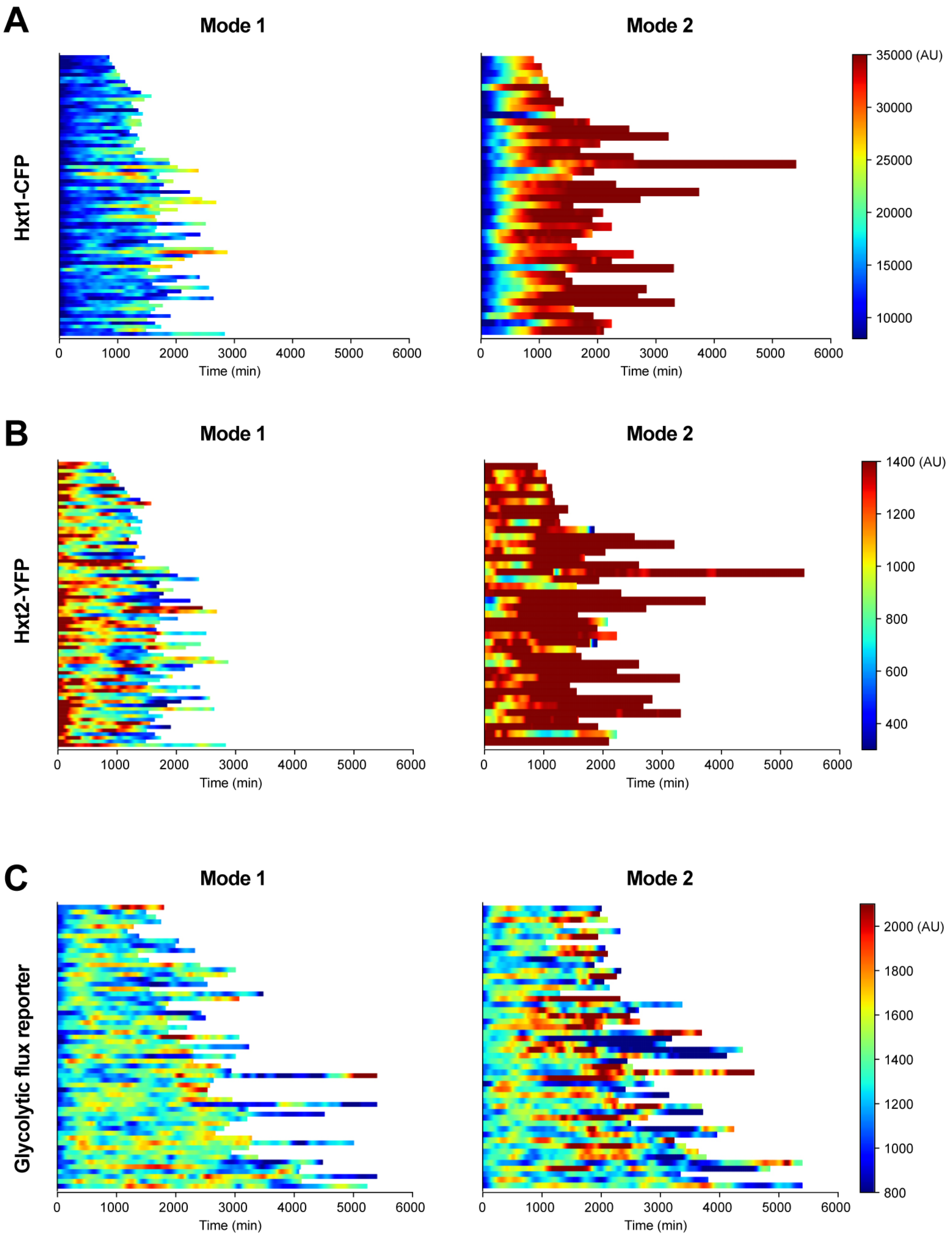


**Fig. S1. Dynamics of various metabolic factors in Mode 1 vs Mode 2 aging.** Single-cell color map trajectories of (**A**) Hxt1-CFP (n=119), (**B**) Hxt2-YFP (n=119), (**C**) glycolytic flux reporter (n=110) in Mode 1 and Mode 2 aging cells. Each row represents the time trace of a single cell throughout its lifespan. Color represents the fluorescence intensity as indicated in the color bar.

A

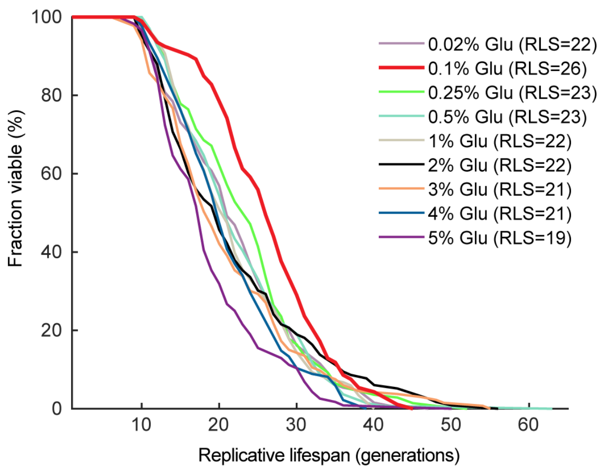


B

**Gehan-Breslow-Wilcoxon test**

| vs. 0.1% Glu | 0.02% Glu | 0.25% Glu | 0.5% Glu | 1% Glu | 2% Glu | 3% Glu | 4% Glu | 5% Glu |
|---|---|---|---|---|---|---|---|---|
| **P-value** | 0.0002 | 0.004 | 0.0002 | <0.0001 | <0.0001 | <0.0001 | <0.0001 | <0.0001 |
| **Significance** | *** | ** | *** | **** | **** | **** | **** | **** |

**Fig. S2. Lifespan curves of WT under different glucose levels.** (**A**) Replicative lifespan curves for WT cells under different glucose levels (0.02% Glucose: n = 130; 0.1% Glucose: n = 93; 0.25% Glucose: n = 140; 0.5% Glucose: n = 136; 1% Glucose: n = 118; 2% Glucose: n = 116; 3% Glucose: n = 133; 4% Glucose: n = 135; 5% Glucose: n = 116). (**B**) Statistical significance of the lifespan differences under various glucose levels. P-values between the lifespan at 0.1% glucose vs lifespans at other glucose levels were calculated by Gehan-Breslow-Wilcoxon test, and the significance is determined as P≤0.01: significant (*), P≤0.001: very significant (**), P≤0.0001: highly significant (***).

A

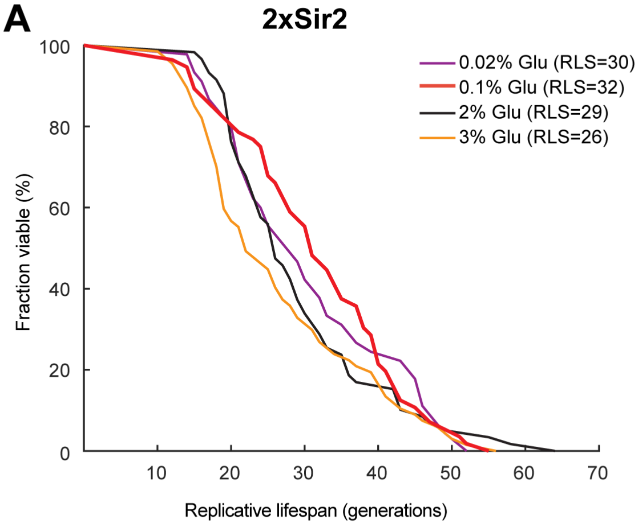


B

**Gehan-Breslow-Wilcoxon test**

| WT vs. 2xSir2 | 0.02% Glu | 0.1% Glu | 2% Glu | 3% Glu |
|---|---|---|---|---|
| **P-value** | 0.0001 | 0.003 | <0.0001 | 0.0017 |
| **Significance** | *** | ** | **** | ** |

**Fig. S3. Lifespan curves of the 2 x *SIR2* strain under different glucose levels.** (**A**) Replicative lifespans for 2x*SIR2* cells under different glucose levels (0.02% Glucose: n = 45; 0.1% Glucose: n = 56; 2% Glucose: n = 59; 3% Glucose: n = 67). (**B**) Statistical significance of the lifespan differences in 2x*SIR2* vs WT under various glucose levels. P-values between the lifespans in 2x*SIR2* vs WT (from Fig. S2) under various glucose levels were calculated by Gehan-Breslow-Wilcoxon test, and the significance is determined as P≤0.01: significant (*), P≤0.001: very significant (**), P≤0.0001: highly significant (***).

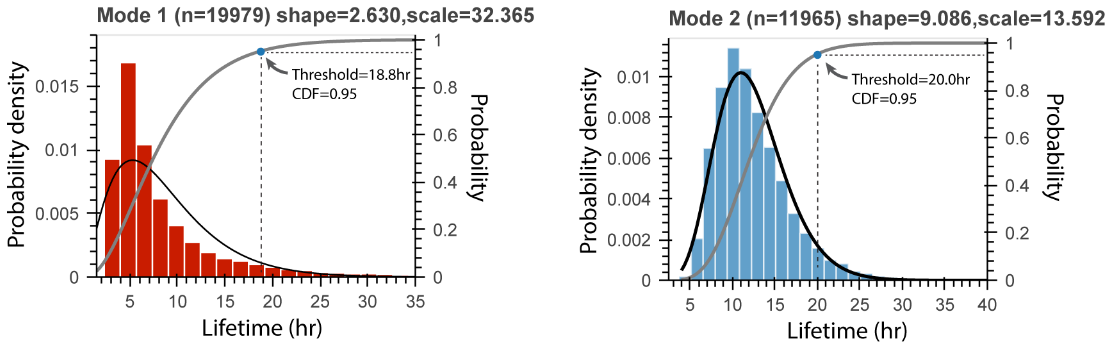


**Fig. S4. Threshold for defining DS cells.** Distributions of simulated lifetime for Mode 1 (red histogram) and Mode 2 (blue histogram) were separately fitted to a gamma distribution with shape and scale shown above the plots. Probability density function (PDF) and cumulative distribution function (CDF) are showed in black and grey lines, respectively. The threshold in each mode for defining DS was set as CDF=95%.

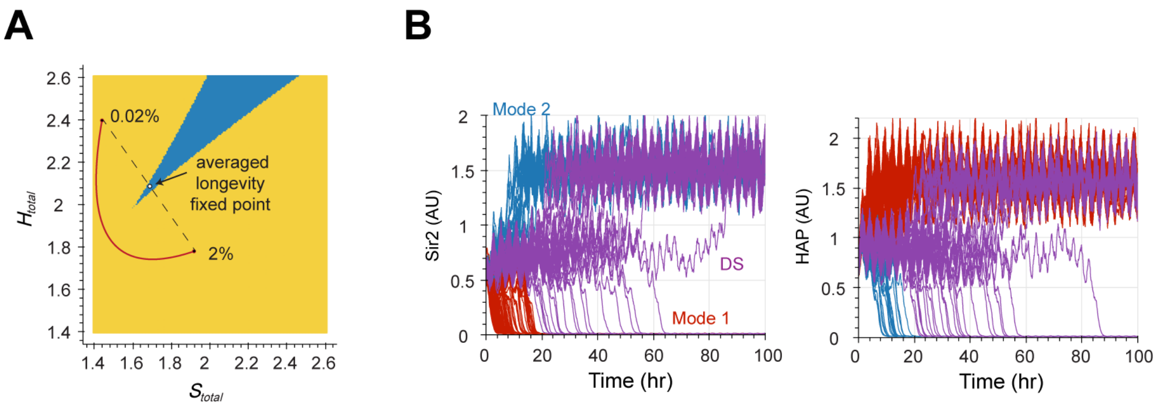


**Fig. S5. Dynamic stabilization does not require the presence of the stable longevity fixed point at an intermediate level of glucose.** (A) The modified path (red curve) of $S_{total}$ and $H_{total}$ corresponding to the glucose levels between 0.02% and 2%, which showed no overlap with the region where the longevity fixed point exists (blue region). The white circle in the blue region represents the averaged stable fixed point generated by oscillations. (B) Stochastic simulations of the Sir2 and HAP time traces during aging under glucose oscillations between 0.02% and 2% from the modified model.

**Movie S1.** Movie for phase plane at glucose concentration changing from 5% to 0.02%. Blue line and red line represent the nullcline of Sir2 and HAP, respectively. Closed circles are stable fixed points and open circles are unstable fixed points. The arrows show the directions of the system's movement.

**Movie S2.** Movie for simulated dynamic behaviors of the system upon external glucose oscillations. The glucose level oscillates between 0.02% and 2% with a period of 6 hours. Left: each purple dot represents a simulated cell at the Sir2-HAP phase plane; Mid: the corresponding simulated time traces of Sir2; Right: the corresponding simulated time traces of HAP.
